# Template-based RNA structure prediction advanced through a blind code competition

**DOI:** 10.64898/2025.12.30.696949

**Authors:** Youhan Lee, Shujun He, Rachael C Kretsch, Toshiyuki Oda, G. John Rao, Yehyun Kim, Raehyun Kim, Hyunjin Kim, Cher Keng Heng, Danny Kowerko, Haowei Li, Hoa Nguyen, Arunodhayan Sampathkumar, Raúl Enrique Gómez, Meng Chen, Atsushi Yoshizawa, Shun Kuraishi, Kenji Ogawa, Shuxian Zou, Alejo Paullier, Bingkang Zhao, Huey-Long Chen, Tsu-An Hsu, Tatsuya Hirano, Wah Chiu, Jeanine G. Gezelle, Daniel Haack, Yibao Hong, Shekhar Jadhav, Deepak Koirala, Anna Lewicka, Shanshan Li, Marco Marcia, Joseph Piccirilli, Boris Rudolfs, Yoshita Srivastava, Anna-Lena Steckelberg, Zhaoming Su, Navtej Toor, Liu Wang, Zi Yang, Kaiming Zhang, Jian Zou, David Baker, Shi-Jie Chen, Maggie Demkin, Andrew Favor, Alissa M. Hummer, Chaitanya K. Joshi, Andriy Kryshtafovych, Emine Küçükbenli, Zhichao Miao, John Moult, Christian Munley, Walter Reade, Theo Viel, Eric Westhof, Sicheng Zhang, Rhiju Das

**Affiliations:** NVIDIA, USA; Texas A&M University, College Station, TX, USA; Biophysics Program, Stanford University, Stanford, CA, USA; QuettaAI, Inc., Tokyo, Japan; Eigen Bio, New York, NY; Professorship of Media Informatics, Chemnitz University of Technology, Chemnitz, Germany; Department of Applied Computer and Biosciences, Mittweida University of Applied Sciences, Mittweida, Germany; CZUR Technology Co., Ltd., Dalian, China; Institut Pasteur, Imaging and Modeling Unit, F-75015 Paris, France; Sichuan Intelligent Expressway Technology Co., Ltd, ChengDu, China; PeptiDream Inc., Kanagawa, Japan; Department of Clinical Engineering, University of Toyama Hospital, Toyama, Japan; Mohamed bin Zayed University of Artificial Intelligence, Abu Dhabi, United Arab Emirates; University of Montevideo, Montevideo, Uruguay; Zhejiang University-University of Edinburgh Institute, Zhejiang University, Haining, Zhejiang, China; Anbogen Therapeutics, Inc., Taiwan, ROC; Division of CryoEM and Bioimaging, SSRL–SLAC National Accelerator Laboratory, Menlo Park, CA, USA; Department of Bioengineering and James Clark Center, Stanford University, Stanford, CA, USA; Department of Microbiology and Immunology, Stanford University, Stanford, CA, USA; Department of Biochemistry and Molecular Biophysics, Columbia University, New York, NY, USA; Department of Biochemistry and Molecular Biophysics, University of California San Diego, CA, USA; Division of Life Sciences and Medicine, University of Science and Technology of China, Hefei, China; Department of Cell and Molecular Biology, Uppsala University, Uppsala, Sweden; European Molecular Biology Laboratory (EMBL), Grenoble, France; Department of Chemistry and Biochemistry, University of Maryland, Baltimore County, MD, USA; Department of Biochemistry and Molecular Biology, The University of Chicago, Chicago, IL, USA; Istituto Italiano di Tecnologia, Genoa, Italy; Science for Life Laboratory, Department of Cell and Molecular Biology, Uppsala University, Uppsala, Sweden; Department of Chemistry, The University of Chicago, Chicago, IL, USA; State Key Laboratory of Biotherapy and Biosafety Laboratory, Institute of Infection and Infectious Diseases, West China Hospital, Sichuan University; The State Key Laboratory of Oral Diseases, National Clinical Research Center for Oral Diseases, National Center for Stomatology, West China Hospital of Stomatology, Sichuan University, Chengdu, China; Department of Cariology and Endodontics, West China Hospital of Stomatology, Sichuan University, Chengdu, China; Department of Molecular Biophysics and Biochemistry, Yale University, CT, USA; Department of Biochemistry, University of Washington, Seattle, WA, USA; Howard Hughes Medical Institute, University of Washington, Seattle, WA USA; Department of Physics and Astronomy, University of Missouri, Columbia, MO, USA; Department of Biochemistry, University of Missouri, Columbia, MO, USA; Institute of Data Sciences and Informatics, University of Missouri, Columbia, MO, USA; Google, Kaggle, Mountain View, CA, USA; Department of Biochemistry, Stanford University School of Medicine, Stanford, CA, USA; Department of Computer Science and Technology, University of Cambridge, UK; University of California, Davis, CA, USA; Guangzhou National Laboratory, Guangzhou International Bio Island, Guangzhou, China; Institute for Bioscience and Biotechnology Research, Rockville, MD, USA; Department of Cell Biology and Molecular Genetics, University of Maryland, MD, USA; Wenzhou Institute, University of Chinese Academy of Sciences, Wenzhou, China; Architecture et Réactivité de l’ARN, Institut de Biologie Moléculaire et Cellulaire du CNRS, Université de Strasbourg, Strasbourg, France; Howard Hughes Medical Institute, Stanford, CA, USA

## Abstract

Automatically predicting RNA 3D structure from sequence remains an unsolved challenge in biology and biotechnology. Here, we describe a Kaggle code competition engaging over 1700 teams and 43 previously unreleased structures to tackle this challenge. The top three submitted algorithms achieved scores within statistical error of the winners of the recent CASP16 competition. Unexpectedly, the top Kaggle strategy involved a pipeline for discovering 3D templates, without the use of deep learning. We integrated this template-modeling pipeline and other Kaggle strategies to develop a single model RNAPro that retrospectively outperformed individual Kaggle models on the same test set. These results suggest a growing importance of template-based modeling in RNA structure prediction.

## Introduction

Computational 3D RNA structure methods have made accurate predictions in blind challenges like RNA-puzzles and the RNA targets of the Critical Assessment of Structure Prediction (CASP)^1–4^. However, successes have typically involved humans ‘in the loop’ of computations, with experts guiding modeling through insights gleaned from published literature, multiple sequence alignments, prior template structures with possible 3D homology, and their own intuition^5–7^. Automated approaches, including a large number of deep learning methods, have underperformed human experts^1–3,8–10^. As a result, despite notable successes, structure prediction has not been broadly adopted for biological inquiry or design of RNA in the same way that it has for proteins.

After observing modest progress by expert groups in the CASP16 RNA structure prediction trials^3,11^, we sought to engage a broader community of non-experts in the RNA structure prediction problem. The Kaggle platform has crowdsourced difficult scientific problems to thousands of participants^12–14^. Compared to CASP and RNA-Puzzles, which engage dozens of expert groups under flexible, assessor-defined evaluation, Kaggle ‘code competitions’ trade off pre-defined metrics and required code upload for broader reach and strict confidentiality of both targets and algorithms.^15^ Recent Kaggle updates allow scoring with evaluation metrics like TM-align^16,17^, which counts the fraction of nucleotides in the predicted and experimental RNA structure that occur within a length-dependent cutoff. The TM-align metric has been calibrated so that scores above 0.45 correspond to correct global folds and has been used to evaluate 3D modeling accuracy in RNA-puzzles and CASP^1–3^.

## Results

### A blind code competition for RNA 3D structure prediction

In February 2025, we launched a Kaggle code competition for 3D RNA structure prediction^18^ (see **Table 1** for complete timeline). The competition was split into two phases. An initial three-month ‘training phase’ solicited algorithms for RNA structure prediction using 23 targets with unreleased structures. A separate evaluation phase was carried out in September 2025. For rigor, the evaluation was based on a completely separate set of ‘future’ 20 targets whose sequences and structures became available in the four months after the close of the training phase in May 2025.

**Table 1.** Timeline of Kaggle RNA Folding Challenge and associated competitions.

| Date | Event |
|---|---|
| May 1, 2024 | CASP16 begins |
| September 29, 2024 | CASP16 last deadline |
| February 27, 2025 | Launch Kaggle competition |
| February 28, 2025 | Parallel RNA-Puzzle launch PZ51, PZ52, PZ54, PZ57, PZ58 |
| March 13, 2025 | Initial MSA release on Kaggle |
| March 21, 2025 | Deadline for parallel RNA-Puzzles PZ51, PZ52, PZ54, PZ57, PZ58 |
| March 26, 2025 | Release scores on Kaggle for AF3 (noMSA/MSA) |
| April 2, 2025 | Release of RibonanzaNet2 (schedule) |
| April 15, 2025 | trRosettaRNA-server updated with end-to-end model (link) |
| April 17, 2025 | Deadline for RNA-Puzzle PZ62 |
| April 23, 2025 | Public leaderboard refresh |
| April 26, 2025 | Early Sharing Prize open (post) |
| May 6, 2025 | BioNemo Youhan Lee posts Boltz-1 notebook (post) |
| May 8, 2025 | KGMON Theo Viel posts ‘Getting Started’ notebook |
| May 10, 2025 | CASP16 RNA assessment bioRxiv posted (link) |
| May 11, 2025 | MSA_v2 released (post) |
| May 13, 2025 | PDB_RNA to enable template discovery (post) |
| May 29, 2025 | Kaggle training phase: Final submission deadline |
| June 23, 2025 | Deadline for RNA-Puzzle PZ63 |
| July 4, 2025 | Deadline for RNA-Puzzles PZ64, PZ65, PZ66 |
| July 29, 2025 | Leaderboard targets run on trRosettaRNA-server |
| August 13, 2025 | Deadline for RNA-Puzzles PZ67 |
| September 5, 2025 | Deadline for RNA-Puzzles PZ68, PZ70 |
| September 24, 2025 | Kaggle competition close (Private leaderboard release) |

The Kaggle training phase was itself split into two periods, with a ‘data refresh’ scheduled between these periods to remove targets that became publicly released in the PDB. During the initial training period of 7 weeks, the 20 RNA targets from CASP16 whose experimental structures were previously unreleased comprised the ‘Public leaderboard’ (**Supplemental Data Table S1**). Participants submitted algorithms that were executed on Kaggle servers, with each algorithm required to generate five 3D predictions per hidden target. The outputs were then scored with TM-align and the mean score across targets was displayed on the continuously updated Public leaderboard (**Fig. 1a**). To lower barriers to entry, a precompiled training data set composed of previously released RNA structures in the Protein Data Bank (PDB) as well as a mock test set based on CASP15 targets^2^ were shared with Kaggle teams. To further lower barriers, competitors’ algorithms needed to only provide a single atom per nucleotide (the C1′ atom, chosen for its centrality), rather than all atoms. In addition, structures were scored based on the best alignment of predicted and experimental nucleotides (TM-align; not TM-score used in CASP scoring, which assumes sequence numbering correspondence), reducing the need for algorithms to optimize nucleotide numbering. Multiple sequence alignments (MSAs) from the field-standard rMSA pipeline^19^ were made available for the training data as well as for the hidden test set sequences. Information on small molecules, proteins, or other nucleic acid partners was also provided, although, for simplicity, there was no requirement to model these interactions, and they were not evaluated. Finally, no restrictions were placed on the use of predictions, scores, or literature from the prior CASP16 competition, although it was noted that such information would not be available for targets in the Kaggle competition’s final evaluation phase. To drive participation, composite scores from two baselines, AlphaFold 3^9^ run with the same rMSA output and a human expert team Vfold^7^, were provided on the competition leaderboard. The Vfold team, which tied for most number of top-RMSD predictions in the last RNA-Puzzles summary^1^ and also led CASP16^3,7^, had a higher baseline score.

**Figure 1.**
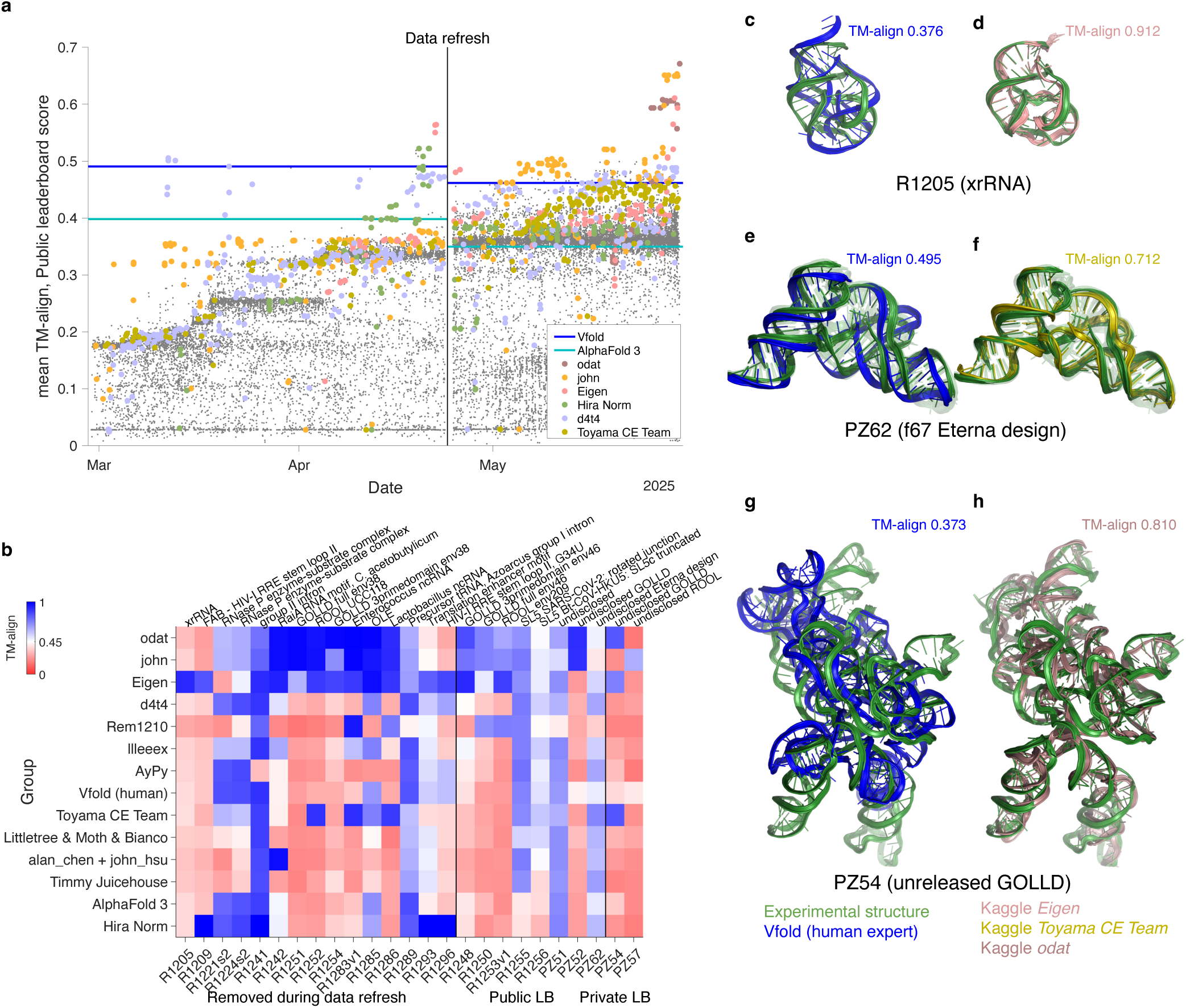
Training phase of the Stanford RNA 3D folding Kaggle competition. **(a)** Public leaderboard (LB) scores (mean TM-align) values of Kaggle submissions vs. time (gray symbols); teams of note are highlighted as colored symbols, and Vfold and AlphaFold 3 baselines are shown as blue and teal lines, respectively. **(b)** TM-align scores for individual RNA targets from top submissions and AlphaFold 3 and Vfold, ordered by mean TM-align scores over the Public leaderboard at the close of the training phase. TM-align > 0.45 (white in heatmap) corresponds to a matching global fold. **(c-h)** Notable blind predictions during the training phase for (**c**,**d**) previously unreleased CASP target R1205, a beet western yellows virus xrRNA (PDB ID: 9CFN^26^); (**e**,**f**) blind RNA-puzzle PZ62, a synthetic design f67 from Eterna (transparent backbones show structural ensemble fit into low resolution cryo-EM map); and (**g**,**h**) blind RNA-puzzle PZ54, monomer from a giant ornate lake- and lactobacillus-derived (GOLLD) RNA^41^ multimer with unreleased sequence and structure. Panels superimpose experimental structures (green) with (**c**,**e**,**g**) best of five Vfold predictions (blue) or (**d**,**f**,**h**) notable Kaggle predictions (from algorithms by Eigen, Toyama CE Team, and odat, respectively). For visualization, all-atom models in (c-h) were derived from C1′-only submissions with Arena^42^.

### Kaggle algorithms close the gap with human experts

Within the first three weeks of the competition, the team ‘d4t4 -l34k -score ’ (hereafter d4t4) was able to outperform both the AlphaFold 3 and Vfold baselines (**Fig. 1a**) by integrating RNA MSA information in Protenix, an open-licensed AlphaFold 3 reproduction^20^. The d4t4 team made publicly available their training algorithm as well as the results of their studies with other codebases and different training sets. Building on these insights and community discussions, other teams, notably Eigen, were also able to achieve scores higher than the Vfold baseline in this first period of the training phase (**Fig. 1a**).

Seven weeks into the training phase, the hidden test set was refreshed to reduce the possibility of the public release of any target data corrupting the rankings. All targets whose structures were anticipated to become publicly available before the end of the training phase were shifted out of the Public leaderboard set. In addition, three RNA-puzzles – concomitantly posed to human expert participants, under conditions of strict confidentiality – were added to the Public leaderboard, giving 8 targets. At the same time, two additional RNA-puzzle targets, PZ54 and PZ57, were included as an initial Private leaderboard, visible only to hosts, to monitor the blind performance of algorithms (**Supplemental Data Table S1**). These targets had structural homologs amongst giant ornate lake- and lactobacillus-derived (GOLLD) and rumen-originating ornate large (ROOL) RNAs that had been recently released in the PDB and had homologs in the training phase Public leaderboard. After the data refresh, the Public leaderboard scores of the Vfold and AlphaFold 3 baselines decreased, and scores of all Kaggle teams decreased below Vfold (**Fig. 1a**), confirming the increased average difficulty of the targets. Interestingly, team Eigen was able to quickly make a submission that restored their leading position in the leaderboard (**Fig. 1a**).

Over the last four weeks of the training phase, other teams began to also produce competitive models, and in the final week, 11 out of 1700 teams assembled from over 10,000 competition entrants submitted algorithms that outperformed Vfold, with two (a late-entering team, odat, and john) outperforming team Eigen. There was a clear gap between the mean TM-align scores for these top three Kaggle teams (0.671, 0.653, 0.615) and the Vfold baseline (0.462) on the Public leaderboard. Furthermore, the top 11 teams did well on different targets, suggesting that they had explored diverse innovations (**Fig. 1b**). For example, for the two Private leaderboard targets, odat, Eigen, and Toyama CE team produced algorithms achieving correct global folds (TM-align > 0.45) for PZ54, while john achieved a good model for PZ57. John, Eigen, and odat all achieved correct global folds for targets homologous to PZ54 and PZ57 in the Public leaderboard, highlighting the importance of the Private leaderboard for differentiating methodological strengths. Subsequent discussions confirmed the diversity of approaches, ranging from the use of RNA foundation models like Aido.RNA^21^ and RibonanzaNet (RNet)^22^ to a wide range of model generation and selection strategies (model descriptions are compiled in **Supplemental Data Table S3**). Unexpectedly, the top two algorithms, from john and odat, explored template-based modeling (TBM) strategies. TBM approaches search for previously solved 3D structures with potential similarity to the targets. While a well-established class of methods^23^, TBM has been largely discontinued in recent RNA modeling pipelines that leverage deep learning^9,20,24,25^. The other top teams were able to retrain AlphaFold reproductions like d4t4’s Protenix code to give improved results without TBM. Eigen’s large boosts came from a specially prepared, up-to-date training set and a high-memory GPU node to handle training examples with lengths greater than 400 nucleotides (**Supplemental Data Table S3**).

**Figs. 1c-h** illustrate the modeling accuracy of the different groups across three targets with different sizes and structures. For target R1205, an Xrn1-resistant RNA from a beet western yellows virus^26^, no CASP16 predictor, including Vfold, achieved the correct fold (TM-align 0.376), while Eigen’s Protenix variant was able to capture the structure accurately (TM-align 0.912; **Fig 1c-d**). **Figs. 1e-f** show a different example, a synthetic target called f67, which was designed on the Eterna platform and had no prior homologous sequence or structure^22^. A blind Vfold prediction submitted to the RNA-puzzle trial for this target (PZ62) captured the secondary structure (the pattern of Watson-Crick-Franklin double helices), and the overall global arrangement of these helices, but the relative positions and orientations of the helices were shifted, leading to a modest TM-align of 0.495 (**Fig. 1e**). Several Kaggle algorithms produced better predictions, with the Protenix variant from the Toyama CE team achieving a TM-align of 0.712, similar to the experimental uncertainty of the structural ensemble fit into the low resolution cryo-EM map (**Fig. 1f**; mean TM-align between ensemble conformations of 0.656).

For both of these targets, Kaggle teams could receive indirect feedback on model accuracy through the Public leaderboard scores. **Figs. 1g-h** shows performance on a fully hidden target on the Private leaderboard, a much larger GOLLD RNA. When faced with this target as an RNA Puzzle (PZ54), human experts on the Vfold team were not able to predict the RNA’s fold (TM-align 0.373; **Fig. 1g**) despite the description of homologous GOLLD structures in two publications^27,28^ and PDB-released entries^29^ during the PZ62 prediction period^27^. In contrast, Kaggle algorithms were able to automatically detect the available template and generate excellent predictions (TM-align > 0.8; **Fig. 1h**).

### Blind evaluation on previously unreleased targets

Automated methods without humans in the loop outperformed the Vfold human expert baseline in the training phase. However, as noted above, for Public leaderboard targets, Kaggle teams benefited from indirect feedback from the composite Public leaderboard scores. Furthermore, for CASP16 targets, the Kaggle teams had an advantage over the Vfold team in making their predictions nearly a year later – the Kaggle algorithms could make use of additional data on targets’ homologs in the PDB as well as published CASP16 structure predictions and scores, even if the targets’ experimental structures were not yet publicly available. To test the predictive power of Kaggle algorithms and to fairly compare performance to Vfold human experts, we therefore collected new targets whose sequences and structures were not released until after the close of the Kaggle training phase, and ran each team’s top two self-selected algorithms on these targets.

The final evaluation set comprised 10 targets that were run as RNA-puzzles for human experts, as well as 10 additional ‘future’ targets released by independent labs in the PDB over the three months subsequent to the Kaggle entry deadline. When run over all 20 targets of this final Private leaderboard, the scores of Kaggle algorithms correlated well with their scores on the Public leaderboard (**Fig. 2a**). The top three teams as ranked by the training phase Public leaderboard (odat, john, and Eigen) were also the top three on this final Private leaderboard (**Fig. 2a**). These three methods each significantly outperformed (*p* < 0.05, one-sided paired t-test) AlphaFold 3 as well as the trRosettaRNA server, which was updated with a performant end-to-end model^8^ at the end of the Kaggle competition (**Fig. 2a** and **Extended Data Fig. 1a**).

**Figure 2.**
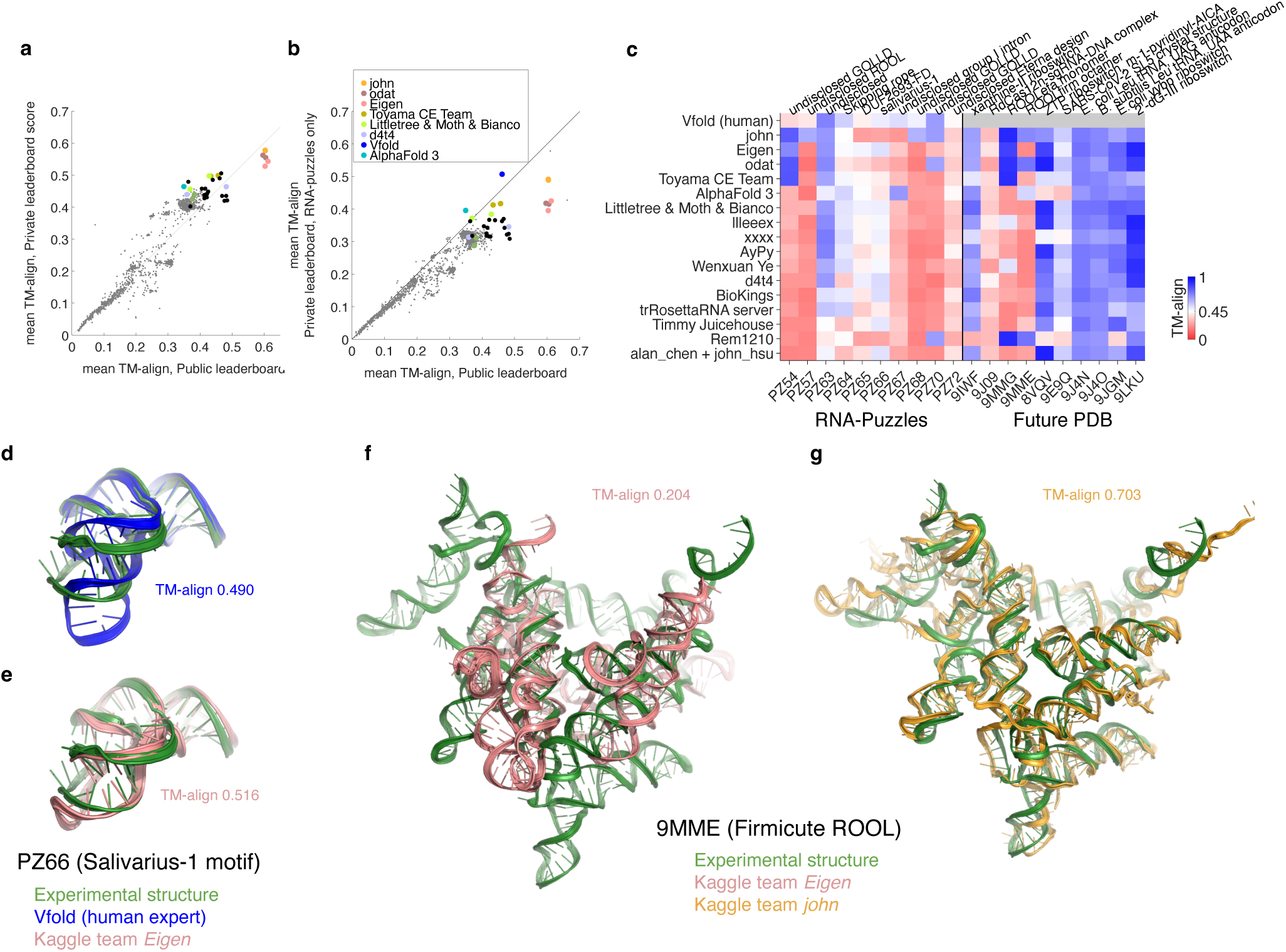
Outcome of final evaluation on Private leaderboard targets. (**a**) Mean TM-align of each Kaggle team’s self-selected top two algorithms in the final Private leaderboard (*n* = 20) vs. Public leaderboard. (**b**) Mean TM-align of Kaggle algorithms and the Vfold group over ten targets that were also made available for human expert prediction as RNA-Puzzles. In (**a**-**b**), black symbols mark top submissions; colors mark specific teams discussed in the main text. (**c**) TM-align scores for individual RNA targets in the final Private leaderboard from top submissions and AlphaFold 3 and Vfold baselines, ordered by mean TM-align over Private leaderboard RNA-Puzzle targets. Some values differ from Fig. 1b, which shows best Public leaderboard submission for Kaggle teams rather than the best of each team’s two self-selected algorithms. (**d**,**e**) Overlay of X-ray crystal structure (PDB: 9G4Q; green) for PZ66 (salivarius-1 motif) and predictions from the Vfold group (blue in **d**) and Kaggle algorithm from Eigen (salmon in **e**). (**f-g**) Overlays of cryo-EM structure for monomer from multimeric 9MME (Firmicute rumen-originating ornate large RNA, or ROOL^43^) with Kaggle predictions from the (**f**) Eigen deep learning model (salmon) and (**g**) template-based modeling pipeline of john (gold). For visualization, all-atom models in (**d-g**) were derived from C1′-only submissions with Arena^42^.

Ten of the 20 Private leaderboard targets were also run as RNA-puzzles for human experts. As has also been seen in protein evaluation^30^, these targets selected for blind challenges were more challenging than the ten structures that arose in the PDB (compare TM-align scores in **Fig. 2a** and **Fig. 2b**; and **Table 2**). Over these more difficult RNA-Puzzles targets, the same three algorithms (john, Eigen, and odat) remained the best Kaggle predictions and achieved mean TM-align scores just below the Vfold team (0.491, 0.425, and 0.418 and compared to 0.507, respectively; **Figs. 2b-c** and **Table 2**). Interestingly, unlike the submissions that ranked best on the Public leaderboard (**Fig. 1b**), john’s selected algorithm for final scoring captured templates for both PZ54 and PZ57 (**Fig. 2c**). Across a variety of ranking metrics (TM-align, TM-score, RMSD; and Z-score-transformed values for these metrics), predictions from john, odat, Eigen, and Vfold achieved top rankings; under lDDT and GDT metrics that are more sensitive to local accuracy and smaller targets (**Extended Data Figs. 1a-c**), Eigen and Vfold generally ranked highest. For the whole Private leaderboard as well as the RNA-Puzzles subset, both john and odat achieved 1st ranking for at least one of the metrics (**Extended Data Figs. 1a-c**). In most rankings and leaderboard subsets, the differences between john, odat, Eigen, and Vfold were not statistically significant (*p* > 0.05, one-sided t-test). While these results suggest near-parity between the Kaggle algorithms and the Vfold expert team, statistical power may be limited by having only 10 RNA-Puzzles.

**Table 2.** Summary of group scores (mean TM-align) over the Kaggle Private leaderboard.

| **Group** | **All<br>(n = 20)** | **RNA-Puzzles<br>(n = 10)** | **Future PDB<br>(n = 10)** |
|---|---|---|---|
| **Baseline** |  |  |  |
| Vfold (human) | undefined | 0.507 | undefined |
| AlphaFold 3 | 0.464 | 0.396 | 0.533 |
| trRosettaRNA server | 0.438 | 0.340 | 0.535 |
| Protenix | 0.409 | 0.304 | 0.515 |
| MMseqs2 | 0.321 | 0.171 | 0.472 |
| Best prior PDB | 0.679 | 0.611 | 0.747 |
| **Kaggle** |  |  |  |
| john | 0.578 | 0.491 | 0.664 |
| odat | 0.562 | 0.418 | 0.706 |
| Eigen | 0.543 | 0.425 | 0.661 |
| llleeex | 0.506 | 0.370 | 0.642 |
| Toyama CE Team | 0.499 | 0.417 | 0.581 |
| Littletree& Moth & Bianco | 0.498 | 0.384 | 0.611 |
| AyPy | 0.485 | 0.366 | 0.605 |
| Wenxuan Ye | 0.479 | 0.363 | 0.596 |
| d4t4 | 0.464 | 0.345 | 0.582 |
| BioKings | 0.458 | 0.343 | 0.573 |
| alan_chen + john_hsu | 0.449 | 0.315 | 0.582 |
| xxxx | 0.442 | 0.367 | 0.516 |
| Timmy Juicehouse | 0.438 | 0.325 | 0.551 |
| Rem1210 | 0.435 | 0.324 | 0.547 |
| Best Kaggle | 0.669 | 0.577 | 0.761 |
| Best blind algorithm | 0.687 | 0.613 | 0.761 |
| **Post-competition** |  |  |  |
| john (TBM only) | 0.591 | 0.483 | 0.700 |
| Agentic Tree Search | 0.630 | 0.549 | 0.711 |
| EBA (TBM only) | 0.576 | 0.488 | 0.664 |
| RNet-3D-EBA | 0.584 | 0.490 | 0.678 |
| RNAPro | 0.648 | 0.541 | 0.755 |
| RNAPro (best public) | 0.636 | 0.516 | 0.757 |
| Best post-competition | 0.684 | 0.597 | 0.772 |

As observed with the Public leaderboard, different methods were effective at different Private leaderboard targets (**Fig. 2c**). For the salivarius-1 RNA, which was also posed to human predictors as an RNA puzzle (PZ66), a blind Vfold prediction recovered the approximate global fold of this RNA (TM-align 0.490; **Fig. 2d**), but with an incorrect bend angle between the molecule’s two coaxial stems. On this target, most Kaggle algorithms fell short of Vfold, but deep learning algorithms like the Eigen model produced more accurate predictions that correctly recovered the bend angle (TM-align 0.516; **Fig. 2e**). For other targets, the TBM-based Kaggle algorithms produced the most accurate predictions. **Fig. 2f** shows the Eigen model for a Firmicute ROOL RNA (PDB: 9MME), which was inaccurate (TM-align 0.204). The TBM algorithms developed by john and odat gave models with higher accuracy (TM-align 0.703 and 0.746, respectively; **Fig. 2g**).

To quantitatively analyze the influence of target and algorithm features across sequence, architecture, context, and experimental methodology (**Supplemental Data Table S2**), we worked with the top-performing Kaggle teams to compile algorithm descriptions (**Supplemental Data Table S3**; other teams produced less accurate predictions on Private leaderboard targets). This analysis confirmed that template availability significantly impacted the performance of the TBM models from john and odat (**Extended Data Figs. 2** **and 3**). Conversely, algorithms that did not use explicit templates, such as Eigen, demonstrated less reliance on templates and instead showed a stronger correlation with sequence length; AlphaFold 3 was also notably sensitive to MSA depth (**Extended Data Fig. 3**). These findings suggested that TBM was optimal for targets with previously available templates, whereas deep learning algorithms were best for short, non-templated targets (e.g., PZ66 in **Fig. 2e**).

**Figure 3.**
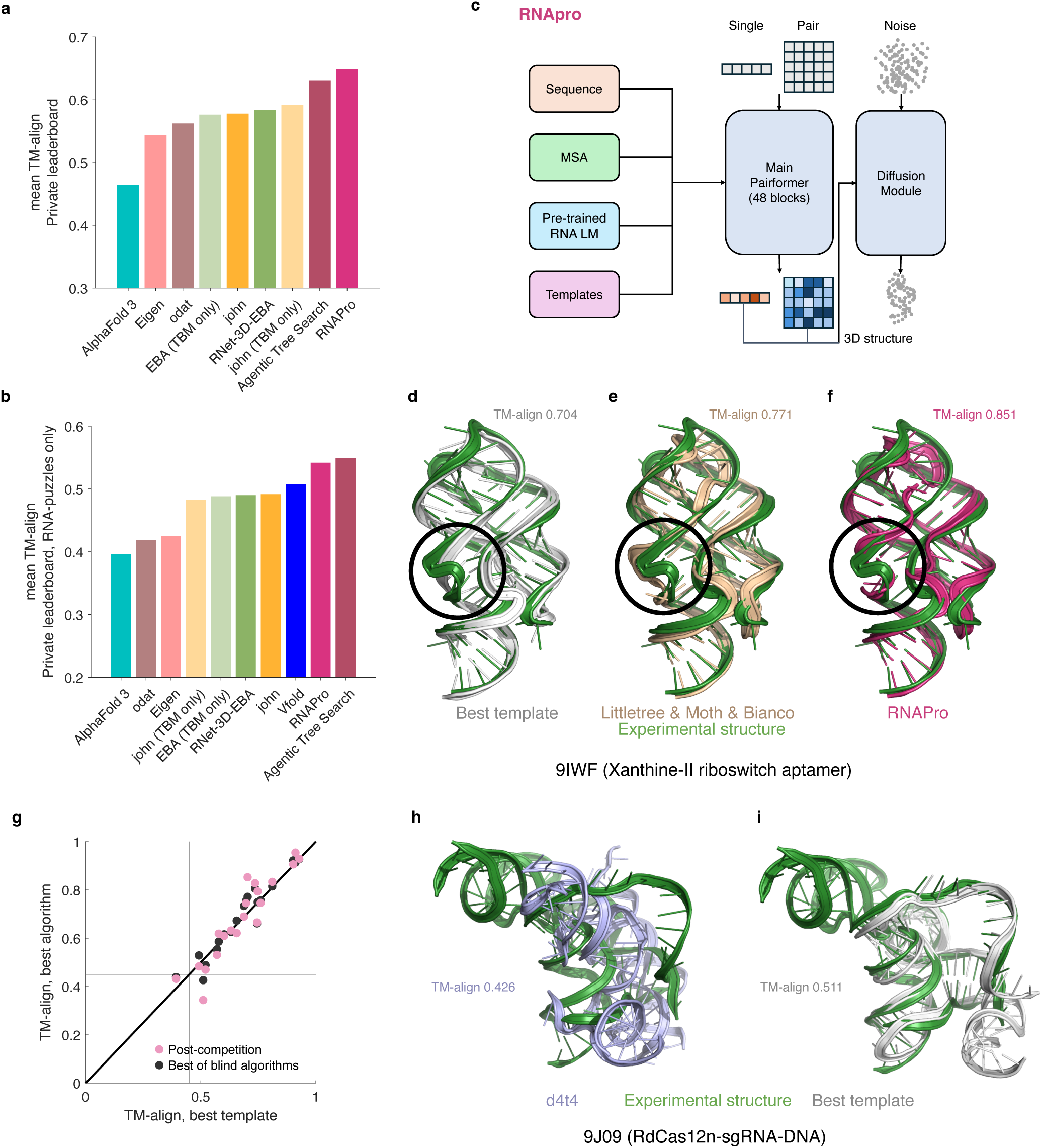
Synthesis of Kaggle models. (a-b) Leaderboard scores of post-competition models over (a) all 20 targets on private leaderboard and (b) just 10 targets that were also RNA-Puzzles, enabling comparison to Vfold human expert. (c) Architecture of RNAPro model. (d-f) Superpositions of the experimental X-ray crystallographic structure of xanthine-II riboswitch aptamer (PDB ID: 9IWF^32^) with (d) best previously available template (PDB ID: 3RKF), (e) deep learning based models from Kaggle algorithm by Littletree & Moth & Bianco, and (f) RNAPro. (g) TM-align modeling accuracies from best of blind algorithms and post-competition models are correlated with highest TM-align templates in the database across the Private leaderboard. Nearly all targets had potential structural templates (TM-align > 0.45; see lines). (h-i) Superpositions of experimental cryo-EM structure of RdCas12n-sgRNA-DNA (PDB ID: 9J09) with (h) best Kaggle model from d4t4 and (i) best prior PDB template (PDB ID: 8EX9). Only sgRNA portions are shown. For visualization, all-atom models in (d-f) and (h-i) were derived from C1′-only submissions with Arena^42^.

### Post-competition synthesis into a single model

The diversity of approaches amongst top Kaggle teams suggested that more accurate models might be achieved by integrating their insights. We paid particular attention to TBM methods since these were important for the top two Kaggle algorithms from odat and john. Indeed, john’s algorithm, which used TBM for some targets and a deep learning method, DRFold2^25^, for other targets, achieved a higher score on all 20 targets of the Private leaderboard when only the TBM method was used (0.578 increasing to 0.591; **Fig. 3a**). The simplest route to integrating Kaggle model insights was to ensemble models produced by distinct approaches. One post-competition model, developed by d4t4 team member arunodhayan through an agentic tree search^31^, achieved a mean TM-align score of 0.630 on the Private leaderboard (**Supplemental Data Table S4** and **Supplemental Data Table S5**). This algorithm ran three top models (Protenix and Boltz implementations by d4t4 and john’s TBM algorithm) and selected five of the fifteen resulting predictions for each target based on a heuristic score that balanced diversity with similarity to the TBM and Protenix models.

In search of a single model that might achieve this performance with less complexity, we developed an updated version of Protenix that integrated MSAs, features from a foundation model RNet2 (see **Methods**), and 3D templates, inspired by innovations from d4t4, Littletree & Moth & Bianco, and john, respectively, and using the updated training data set from Eigen (**Supplemental Tables S2-S3**). The resulting model, called RNAPro (abbreviation for ‘RNA Protenix’, **Fig. 3c**), outscored Kaggle models in the Private leaderboard score (mean TM-align 0.648), and ablation studies confirmed that templates were the most important RNAPro input (**Extended Data Fig. 4a**). We note that a prospective blind evaluation is required to confirm generalization beyond this retrospective analysis, and further development will be required for RNAPro to emulate baseline models with respect to all-atom metrics and local accuracy (**Extended Data Fig. 5**). In particular, for targets with partial templates, RNAPro can collapse C1′ atoms in regions missing from the templates, necessitating post-processing or repeated re-sampling for downstream applications. Nevertheless, for some targets, like the xanthine-II riboswitch aptamer (9IWF^32^; **Figs. 3d-f**), RNAPro gives better predictions (TM-align 0.851; **Fig. 3f**) than any of the Kaggle algorithms (Littletree & Moth & Bianco, TM-align 0.771; **Fig. 3e**), resulting in a higher TM-align score than the best prior template in the PDB (guanine riboswitch 3RKF, TM-align 0.704; **Fig. 3d** and **Supplemental Data Table S6**), despite not explicitly modeling the xanthine ligand bound to the target. The main improvements were in irregular loops interconnecting the three helices of this RNA (circles in **Figs. 3d-f**). Although these results were obtained retrospectively, they suggest a potential synergy between TBM and deep learning refinement. Nevertheless, as in CASP16 RNA structure prediction^3^, the accuracy of models from these post-competition efforts remained strongly correlated with the highest similarity of previously solved structures to the target (**Fig. 3g** and **Extended Data Fig. 3**).

Interestingly, changing the number of training steps or the strategy for template search led to increased accuracy on some targets but lowered accuracy on others (**Extended Data Fig. 4a-b** and **Supplemental Data Table S5**). In addition, use of an embedding-based analysis (EBA^33^) for template discovery, inspired by the Kaggle algorithm from odat, was important for PZ67, an undisclosed group I intron, but missed templates in other cases (**Fig. 3a-b** and **Table 2**; see also **Extended Data Fig. 4b** and **Supplemental Data Table S5**). These results suggest that further improvements might be achievable with further exploration. Indeed, for one target 9J09 (an RdCas12n-sgRNA-DNA complex^34^), all tested modeling approaches gave worse accuracy (best model TM-align 0.426 from d4t4; **Fig. 3h**) than a partially homologous template available in 8EX9 (ISDra2 TnpB in complex with reRNA and cognate DNA^35^, TM-align 0.511; **Fig. 3i** and **Supplemental Data Table S6**). The similarity of the target and template molecules’ functions – TnpB is an evolutionary ancestor of Cas12 – hints that further improvement in template search may come from automated integration of functional annotations and scientific publications^36^, which was not explored in Kaggle models (**Supplemental Data Table S3**).

These results suggest a growing importance of TBM in RNA structure prediction. In this Kaggle competition (2025), 19 of 20 total targets (95%) had potential templates, defined as prior PDB structures with TM-align greater than 0.45 (**Fig. 3g** and **Supplemental Table S6**). In contrast, in CASP15 (2022), only 3 of 12 (25%) targets had potential templates with an increase to 56% of templated targets (19 of 34) in CASP16 (2024).^3^ For example, the large GOLLD and ROOL RNAs, which were intractable in CASP16 due to their novelty, were solvable in this Kaggle competition due to the intervening publication of representative structures and the newer template search methods developed here. Still, these and other Kaggle targets would be classified in CASP as ‘TBM-hard’ or requiring free modeling (‘TBM/FM’)^37^ – neither Vfold nor any single algorithm successfully leveraged templates for all of them. Results for other RNA-puzzles prediction teams are unavailable until the next RNA-Puzzles summary paper, but analysis of those predictions will reveal if other experts or more sophisticated sequence-based algorithms^38–40^ were able to detect all templates. Generally, the recent growth in the experimental database of complex RNA structures, driven by accelerations in RNA structure determination methods, is expected to increase the importance of templates.

## Discussion

We have described a Kaggle machine learning competition that has recruited thousands of people to tackle the automated prediction of 3D RNA structure from sequence. Engaged by a leaderboard providing feedback on hidden targets, the top teams developed algorithms that reduced the gap with CASP16 winners Vfold on confidential targets. TBM was a key feature of the two best entries, and the integration of TBM with deep learning in a post-competition model RNAPro suggested a route to further improvements in accuracy. Because most targets had potential templates, comparative conclusions favoring TBM may not generalize to template-poor RNAs; it remains to be benchmarked whether template-independent algorithms, like team Eigen’s method and AlphaFold 3, might rival RNAPro in the template-free regime. Furthermore, this competition did not evaluate all-atom stereochemistry, local accuracy, large synthetic RNA molecules, the recovery of noncanonical pairs, confidence estimation, modeling of RNAs with multiple 3D structures, RNA with chemical modifications, small molecule binding by RNA, nucleic acid-nucleic acid interfaces, or nucleic-acid protein interfaces, all of which have been challenging for blind prediction^2,3,10^ and are important future directions for RNAPro. As the dominant determinant of accuracy in this benchmark was template availability rather than length or motif complexity (**Extended Data Figs. 2-3**), focused experimental determination of RNAs without prior templates may be a particularly impactful investment for the field, both to expand the knowledge base for template-based modeling and to evaluate progress in template-free modeling. As the field strives for these next milestones, further blind code competitions that reach out to large non-expert communities may be useful to drive progress.

## Acknowledgements

This paper is dedicated to the memory of Neocles Leontis. We thank Elizabeth Park and Addison Howard (Kaggle) for administration of the Kaggle competition; Wei Huang (Case Western) for contributing CASP16 targets to seed the Public leaderboard; Stanford-SLAC Cryo-EM Center, U.C. Berkeley, ESRF CM01, and EMBL Heidelberg cryo-EM facilities for assistance and support in cryo-EM data collection; Advanced Photon Source, a U.S. Department of Energy (DOE) Office of Science User Facility operated for the DOE Office of Science by Argonne National Laboratory under Contract No. DE-AC02-06CH11357, for crystallography support; Hamish Blair, Ann Kladwang (Stanford), the NVIDIA DGX Cloud team and NAIRR Pilot (allocation NAIRR240281) for enabling release of RNet2 alpha during the competition; and the NHR high-performance computing facilities at TU Dresden (ZIH), available through the National High-Performance Computing Alliance for computing support.

## Author contributions

S.H., Y.L., D.B., M.D., A.F., A.M.H., A.K., E.K., R.C.K., Z.M., J.M., C.M., W.R., T.V., E.W., and R.D. developed and hosted the Kaggle competition. T.O., G.J.R., Y.K., R.K., H.K., C.K.H., D.Kow., H.L., H.N., A.S., R.E.G., M.C., A.Y., S.K., K.O., A.P., B.Z., S.Zo., H.-.L.C., T.-.A.H., and T.H. developed and documented Kaggle algorithms. J.G.G., W.C., D.H., Y.H., S.J., D.Koi., R.C.K., A.L., S.L., M.M., J.P., B.R., Y.S., A.-.L.S., Z.S., N.T., L.W., Z.Y., K.Z., and J.Z. provided experimental target structures. S.-.J.C. and S.Zh. provided Vfold predictions. Y.L., S.H., A.S., D.Kow., C.K.J., C.M., and T.V. developed post-competition models. R.D. wrote the manuscript with input from all authors.

## Methods

### Competition setup

Training sequences and structures provided to Kaggle participants were collated from the PDB with custom scripts available at https://github.com/Shujun-He/Stanford3Dfolding_dataprocessing. Multiple sequence alignments for training and test sequences were generated with the rMSA pipeline^19^. Test sequences were collated from CASP16 targets and from unreleased x-ray crystallographic and cryo-EM structures from the labs of the competition hosts and collaborators. The non-CASP targets were run as RNA-puzzles, which required human expert participants to keep sequences strictly confidential and to agree ahead of time not to participate in the Kaggle competition. Solution files, baseline submission files, and scoring was carried out with Python scripts available at https://github.com/DasLab/k1_tools. The Vfold team agreed to share blind predictions to provide baseline scores for this Kaggle competition (see below).

As described in the main text, the Public leaderboard was refreshed in the middle of the training phase to remove targets that were expected to be publicly released by the end of the competition. For the final Private leaderboard, thirteen targets were collected from entries released in the PDB after the close of the training phase. These were manually curated to be composed mainly of RNA. Seven of these PDB targets had no homology to prior PDB structures identifiable by MMseqs2^44^, the PDB’s sequence-based search tool. Three of these targets (9G4P, 9G4R, 9G4Q) were also RNA-Puzzles (PZ64, PZ65, PZ66) and are listed as such throughout the manuscript.

Three targets (9E9Q, 9J4N, 9J4O) had MMseqs2-identifiable templates, but visual inspection by the structure authors noted conformational differences compared to prior structures. Another three targets (9MMG, 8VQV, and 9LKU) had near-sequence identical homologs and were included as positive controls. See **Supplemental Data Table S1** for detailed descriptions of targets.

A text file in comma-separate values format called ‘test_sequences.csv’ held the target information. This file included for each target an anonymized target id; the RNA sequence; a ‘temporal cutoff’ corresponding to the target release date or, for CASP or RNA-Puzzle targets, the target’s deadline in those competitions; short natural language descriptions of the molecule, as are made available in CASP; and a list of all protein and nucleic acid sequences and small molecules in the solved complex in FASTA format. **Supplemental Data Table S1** provides an example. To help competitors during training, an example of this file, curated for CASP15 targets, was provided with the competition data, but the actual test sequences file used for algorithm scoring was kept fully hidden and was only viewable by algorithms once submitted to the Kaggle server. This arrangement kept both the sequences and structures of the targets fully confidential.

Multiple sequence alignments were prepared with rMSA^19^, as follows: rMSA was downloaded from GitHub (https://github.com/pylelab/rMSA) and run using default settings. To convert the rMSA outputs, in which RNA sequences are formatted as DNA sequences, all "T" nucleotides were replaced with "U". The MSAs made available to Kaggle teams during the competition phase were created with the following database versions, released in or before 2022 to correspond as closely as possible to the temporal cutoff for the CASP15 validation set (2022-05-28) suggested to the competitors: RNAcentral^45^ (v20.0, 2022-03-28), RFam^46^ (v14.7, 2021-12-09), and NCBI Nucleotide (2022-10-03). The MSAs used as input for the AlphaFold 3 baseline models and for the final evaluation set were generated with the most up-to-date databases available at the close of the training phase: RNAcentral (v25.0, 2025-05-14), RFam (v15.0, 2024-09-10), and NCBI Nucleotide (2025-06-17).

A few targets had multiple possible conformations, arising from cryo-EM 3D classification, coordinate uncertainties from auto-DRRAFTER^47^, alternative conformers in the asymmetric unit of a crystal structure, or multiple available crystal structures with small sequence variants. In these cases, all structures were recorded as possible ground-truth coordinates (see next).

Examples of these ground truth sequences and C1′ coordinate ‘labels’ were provided to competitors in validation files with the 12 RNA targets from CASP15. As further auxiliary data for the competition, an updated version of the RNet foundation model (called RNet2 alpha), trained on chemical mapping data for 40 million sequences, as well as 400,000 RNA synthetic structures developed with RFDpoly^48^, an updated version of the RFdiffusion design code, were released, but these were not used in top solutions.

Each algorithm developed by Kaggle participants was submitted as a Python notebook that was run on Kaggle servers with access to any data sets associated with the notebook and to a confidential version of the competition data folder with the hidden test sequences and their respective rMSA outputs, but without access to the internet. Submission frequencies were limited to two notebooks per team per day. Notebooks were required to produce C1′ coordinates for five models for each target. Descriptions of notable Kaggle models are provided in **Supplemental Data Table S3**.

### Baseline models

Five predictions for human expert team Vfold for CASP targets were downloaded from the CASP16 website. All other Vfold predictions were submitted in the context of RNA-Puzzles blind predictions (**Table 1**) and sets of five models per target were made confidentially available to Kaggle competition hosts as baselines. The Vfold structure prediction method included three components: prediction of the secondary structure, identification of 3D templates from known structures, and coarse-grained molecular dynamics (CGMD) simulations guided by the secondary structure and templates. Template identification was based on the motifs (e.g., hairpins, junctions, bulges). However, templates often could not be found. In such cases, Vfold either adopted templates of lower sequence similarity or ran coarse grained MD simulations without using templates. Vfold failed to identify ideal templates for PZ54 and PZ57 due to the lack of available GOLLD and ROOL RNA structures in the template database, which was not updated until after those puzzles. However, in those cases, templates for certain local structural elements, such as hairpin loops, were successfully identified and used in the structure prediction; global templates for later test cases involving GOLLD and ROOL structures were identified.

AlphaFold 3^9^ was run as follows: AlphaFold 3 was installed locally and run using JSON file input. Models were generated with AlphaFold 3 version 1, and one seed (seed = 1). AlphaFold 3 was run with the Kaggle competition rMSA input by setting the "unpairedMsa" parameter to the MSA generated by rMSA; this is the AlphaFold 3 model in paper figures. As an additional baseline, AlphaFold 3 was run without any MSA input by setting the "unpairedMsa" parameter to an empty string; these scores were lower than runs with the MSA input. trRosettaRNA server models were obtained from the online server with default settings (https://yanglab.qd.sdu.edu.cn/trRosettaRNA/); the code for the trRosettaRNA2 package, updated after the CASP16 competition^8^, remains unpublished. To avoid leakage, sequences were run on the server on July 26, 2025, after the close of the Kaggle training phase. The trRosettaRNA server returned single models for each sequence; its rankings relative to other methods remained similar (e.g., consistently lower than top Kaggle algorithms and Vfold) when other methods were similarly evaluated based on their first rather than all five models (**Supplemental Table S5**). Protenix (commit 205b65e) was downloaded and run from the code available at the time of the competition close: (https://github.com/bytedance/Protenix/commit/205b65e7223e5ae8dcb9261b78066757bf879c27)

As an additional baseline, a simple template-based modeling workflow using MMseqs2 was made available to Kaggle predictors as a baseline, using the PDB files provided as Kaggle competition data; see https://www.kaggle.com/code/rhijudas/mmseqs2-3d-rna-template-identification.

As a final baseline, USalign (see next section) was used to discover the best possible 3D template available during the competition by aligning the experimental structure against all PDB RNA chains published before the training phase close date, May 29, 2025.

### Model scoring

Outputs of C1′ coordinates from Kaggle notebooks were scored against ground truth experimental coordinates with a publicly available evaluation notebook (https://www.kaggle.com/code/metric/ribonanza-tm-score?scriptVersionId=224830487). The evaluation notebook extracted in PDB format only the target sequence, a single RNA chain, from the model and a ‘solution’ file and computed the accuracies of models compared to experimental coordinates as follows:

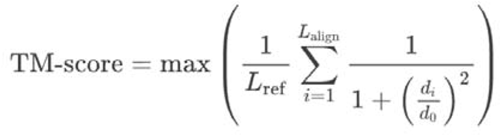

where:

● *L*_ref_ was the number of residues solved in the experimental reference structure (“ground truth”).
● *L*_align_ was the number of aligned residues.
● *d*_i_ is the distance between the i_th_ pair of aligned residues, in Angstroms.
● *d*_0_ is a distance scaling factor in Angstroms, defined as:

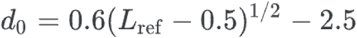

for L_ref_ ≥ 30; and *d*_0_ = 0.3, 0.4, 0.5, 0.6, or 0.7 for L_ref_ <12, 12-15, 16-19, 20-23, or 24-29, respectively. These length-dependent distance scaling factors have been normalized so that TM-scores above 0.45 reflect true fold similarity, irrespective of structure length^16^. TM-scores are computed over one representative atom per residue, here chosen to be C1′. The scoring was carried out with US-align^17^, downloaded from https://github.com/pylelab/USalign on February 22, 2025, run with the flag -atom " C1′". These settings led to TM-scores being computed after superposition of the model and experimental structure and aligning residues independent of their sequence, a mode referred to as TM-align in CASP RNA assessments and in this manuscript. The best TM-align computed across the five models for each target scored against each experimental structure for the target was then averaged across the Public or Private leaderboard targets to give the respective leaderboard scores. As in other Kaggle competitions, teams were allowed to select two submitted algorithms during the training phase as final submissions for ranking on the Private leaderboard. For any team that did not make the selections, the two algorithms that the team submitted during the training phase with the highest Public leaderboard scores were automatically selected as the team’s submissions.

After the competition, rankings were revisited with USalign run with the flag “-TMscore 1”, which forces sequence correspondence between each prediction and experimental structure, called TM-score values in CASP analyses. C1′ RMSD values were calculated in Python with the Kabsch algorithm^49^. GDT values were computed with the TMscore executable in USalign; lDDT values were computed with OpenStructure^50^, with C1′-only reference models. For each of these metrics, Z-scores were computed based on the number of standard deviations of scores away from the mean of the scores for each target; the standard deviation and mean values were recomputed after removing outliers with Z scores less than –2 in a first round, and Z-scores were clipped to be greater than or equal to zero^2,3^. A composite Z-score was defined as Z_composite_ = 0.3 Z_TM-score_ + 0.3 Z_GDT_ + 0.4 Z_lDDT_; weights for this composite metric were analogous to the CASP16 composite Z-score^3^, but individual metrics were calculated here with C1′-only rather than all atom models.

In addition to C1′-only analysis, all-atom predictions for RNAPro, Protenix, AlphaFold (with rMSA and no MSA), and Vfold were analyzed. The O2′ atoms from the RNAPro models were removed and rebuilt using Arena^42^. All-atom models were derived from the C1′-only models submitted by teams john, odat, and Eigen were built using Arena. Any residue in the experimental structure that was non-canonical or different in sequence to the predicted sequence was ignored. Any predicted model that had the incorrect sequence was ignored (**Supplemental Table S7**). The following metrics were calculated: (1) TM-align and TM-score, using USalign with default C3′ atoms or C1′, (2) GDT-TS using LGA using the default C4′ and C1′, (3) lDDT using OpenStructure using the default all-heavy-atom and C1′-only and performing the default stereochemical checks and not, and (4) RMSD using Kabsch algorithm for the default all-heavy-atom and C1′ atoms. Models that had C1′ or C4′ atoms very close or overlapping errored in the LGA run and were hence given a GDT-TS score of 0 (**Supplemental Table S7**). Models where no residue passed the OpenStructure default stereochemical checks were assigned lDDT of 0 (**Supplemental Table S7**). For each target, these scores were converted to Z-scores as above, and then the sum of Z-scores greater than zero was calculated for each group. A composite Z-score developed in CASP16 was also calculated as a summary score: Z_CASP16_ = 0.3 Z_TM-score_ + 0.3 Z_GDT_ + 0.4 Z_lDDT_.

### Model visualization

To aid 3D visualization, all-atom models were derived from the C1′-only Kaggle submissions with Arena^42^ and visualized in Pymol (https://www.pymol.org/); these all-atom models were not used for Kaggle competition scoring.

### Target labeling

To help interpret the strengths and weaknesses of different algorithms across different targets, targets were labeled based on sequence and structure features. Sequence descriptors, sequence length and GC-content, were calculated based on the sequence supplied to predictors. The normalized number of effective sequences (Neff) quantifies the number of non-redundant sequences, here defined as sequence identity less than 80%, in the MSA files provided to Kaggle predictors, based on rMSA (see above). The sequence identity between each pair of sequences in an MSA was calculated after ignoring any positions that were gapped in both of the sequences. For each sequence in an MSA, the weight for that sequence was calculated as the number of other sequences in the MSA with a sequence identity above 80%. The Neff value was then calculated as the sum of inverse weights for each sequence in the MSAs. The template availability was quantified by the TM-align of the closest template as described in the Baseline section. The radius of gyration was calculated as 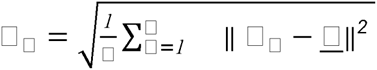 with □□ as the position of atom □, □ as the mean position of all atoms, and || ||^2^ as the squared Euclidean length of a vector. The radius of gyration was normalized by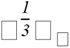 , with □ the number of nucleotides modeled and □□ as monomer length of a coil model (3.4 ); the exponent of ⅓ was selected to represent maximal compaction. Base pair and motif annotation were obtained from Rosetta’s rna_motif^51^. Base pairs were labeled as canonical if they were GC, AU, or GU base pairs involving the Watson-Crick edges of each base; all other base pairs were labeled non-canonical. These base pairs were used to calculate the fraction of bases that were paired as well as the percentage of base pairs that were canonical. Canonical base pair labels were used to determine the number of multi-way junctions, defined as loops where more than 2 stems terminated. The experimental method (x-ray or EM), context (ligand, homo-oligomer) and functional type were manually labeled. These features can be found in **Supplemental Table S2**.

### Post-competition Kaggle synthesis

An automated agentic tree search^31^ guided by Private leaderboard scoring after the competition close was used to develop a model integrating the complementary strengths of notebooks running TBM, Protenix, and Boltz. The weighted agentic tree-search ensemble selected the most reliable five conformations from the 15 candidate structures produced by the three models (5 per model). For each RNA target, the ensemble was anchored with two strong priors – Protenix conformation 2 and TBM conformation 1 – which consistently provided stable local geometry and template-guided global topology. The remaining conformations were evaluated iteratively through a greedy agentic search that balanced three criteria: (i) model reliability, using weights that prioritized Protenix (0.75) over TBM (0.15) and Boltz (0.10); (ii) structural diversity, quantified as the mean pairwise Kabsch-aligned RMSD between the candidate and the selected structures (*diversity*); and (iii) geometric consistency with priors, penalizing excessive deviation from the anchor conformations (*distance_to_priors*). These terms were combined into a weighted scoring function *score* = *w*_model_ ( *w*_div_ *diversity* - w_dist_ *distance_to_priors* ), where *w*_model_ encoded the model-specific reliability weight, *w*_div_ controlled the reward for diversity, and *w*_dist_ controlled the penalty for drifting from the priors. At each iteration, the agent selected the conformation with the highest score, repeating the process until five conformations were chosen.

In an effort parallel to the Kaggle competition, a structure prediction model named RNet-3D was built by fine-tuning the RNet (RibonanzaNet)^22^ foundation model, pre-trained with chemical mapping profiles, to predict 3D coordinates without auxiliary data. Inspired by the findings of Kaggle team odat, Rnet-3D-EBA was further developed after the competition to leverage templates discovered by embedding based alignment (EBA)^33^. RNet-3D largely followed AlphaFold 3^9^, with a few distinctions. First, it did not use triangle attention, to save memory and compute. Second, it did not include multiple sequence alignments as inputs to avoid complexity during training. Third, instead of inputting real templates during training, masked template modeling was used. In this procedure, incomplete templates (masked by up to 50%) were generated from ground truth C1′ coordinates. (During inference, real templates by any template generation process can be used.) The decision to use masked template modeling was largely due to sparsity of RNA structural data, in which most training sequences did not have good templates. All distograms were computed with C1′ coordinates and clipped at 39 Å. RNet2 embeddings (https://www.kaggle.com/models/shujun717/ribonanzanet2) and template embeddings from a template PairFormer were used as inputs to a PairFormer module to generate conditioning for non-equivariant diffusion. RNet-3D was trained for 48 epochs with learning rate starting at 0.0005 and annealed to 0 from epoch 22 to 48, with a batch size of 8 and diffusion module batch size of 48. Masked template modeling was used 50% of time during training; otherwise, the model was trained to make predictions from RNet2 embeddings alone. PDB entries up to March 1, 2024 were used for training. During inference, templates were generated with embedding based alignment against the PDB structures up to May 29, 2025. The Smith-Waterman gap penalty was set to -4 and open gap penalty to of -10, and sum of row/column normalized Z-scores of exponentiated L2 distances of embeddings were used as similarity scores. PDB structures were ranked based on cosine and L2 distance to the target sequence, and Smith-Waterman alignment was carried out for the top 500 most similar structures to find templates. The code for template search is available at https://www.kaggle.com/code/shujun717/rnet2-alpha-template-search-public. For RNet-3D inference, 10 recycling runs and 200 diffusion steps were used to create the 3D structures. For each target, 5 predictions were made, where the first 4 predictions were made with top 4 templates based on Smith-Waterman alignment score, and the last prediction was made with only RNet2 embeddings as inputs to cover targets without templates. The full inference code is available at https://www.kaggle.com/code/shujun717/rnet-3d. While this model outperformed individual Kaggle models in the Public and Private leaderboard, it was less performant than RNAPro, described next.

As a more fully integrative approach to synthesizing Kaggle insights, an end-to-end (E2E) model, called RNAPro, was designed and trained. This model integrated template modeling, multiple sequence alignment (MSA), and a pre-trained RNA language model using the end-to-end training and inference in Protenix (commit 205b65e). During training, for MSA modeling, the MSA module utilized was identical to that used in Protenix. For template modeling, a total of 45 templates were used to obtain diverse templates: 40 obtained from MMseqs and 5 derived from john’s first place Kaggle solution. When no suitable template was found, a zero-valued template was used. During training, 4 templates were randomly sampled from the 45 available templates. The distogram features extracted from C1′ templates were added to the pair representations, which were subsequently updated using the Pairformer module for both single and pair representations.The pre-trained RNA language model RNet2 (https://www.kaggle.com/models/shujun717/ribonanzanet2) was utilized. Single and pairwise representations from each layer of RNet2 were combined via weighted averaging and employed to update the respective representations in the model. Gating modules using a sigmoid function were utilized for these updates. The model parameters were initialized from Protenix base model checkpoints, while new modules, such as the template encoder, were newly initialized. The RNA foundation model (RNet2) remained frozen during training, whereas parameters in the gating module were updated. The training of RNAPro was conducted in two stages. In the first stage, the model was trained for 50,000 steps using the Kaggle dataset, which included PDB entries up to Dec. 17, 2024. In the second stage, a dataset of up-to-date sequences and MSAs made publicly available by team Eigen (cutoff extended to May 29, 2025) was added to the Kaggle data, and training was continued for another 50,000 steps. The learning rate was 0.0005 and 0.0001 for the first and second training, respectively. The diffusion batch size was set to 8, and the training crop size was 512, with random cropping applied during all training. The loss function and its weights were identical to those used in Protenix and included no explicit non-bonded clash or steric repulsion term. Due to the large memory requirements of the integrated model, eight high-memory-enabled Blackwell GPUs (GB300) were used during training. For inference, template sampling was varied to generate diverse structural predictions. Five templates were selected using the template discovery method developed in john’s Kaggle-winning algorithm (https://www.kaggle.com/code/jaejohn/rna-3d-folds-tbm-only-approach), which used templates extended from the competition data training examples up to May 7, 2025. Top 1 through top 5 templates were used to generate five distinct predicted structures. To accelerate computation, cuEquivariance (https://github.com/NVIDIA/cuEquivariance) was employed.

## Usage of AI tools

Claude (Opus 5.0) was used during preparation of the revised manuscript for editorial assistance and preparation of Source Data files; all AI-assisted output was reviewed and verified by the authors.

## Data Availability

Kaggle competition training data are available at https://www.kaggle.com/competitions/stanford-rna-3d-folding/data. Data were processed with scripts available at: https://github.com/Shujun-He/Stanford3Dfolding_dataprocessing. Scripts for scoring and solution compilation are available at: https://github.com/DasLab/k1_tools. Scores for post-competition algorithms can be confidentially computed through submission of algorithms at the Kaggle competition. Solution files for the competition will be released upon publication of the target structures.

## Code Availability

All Kaggle algorithms described herein have been made publicly available as open source notebooks with documentation through ‘Solution’ links at https://www.kaggle.com/competitions/stanford-rna-3d-folding/leaderboard, and links in Supplemental Table Files S2-S3. RNAPro code is available on Github (https://github.com/NVIDIA-BioNeMo/RNAPro), with weights at https://catalog.ngc.nvidia.com/orgs/nvidia/teams/clara/collections/rnapro/artifacts.

---

**Extended Data Fig. 1.**
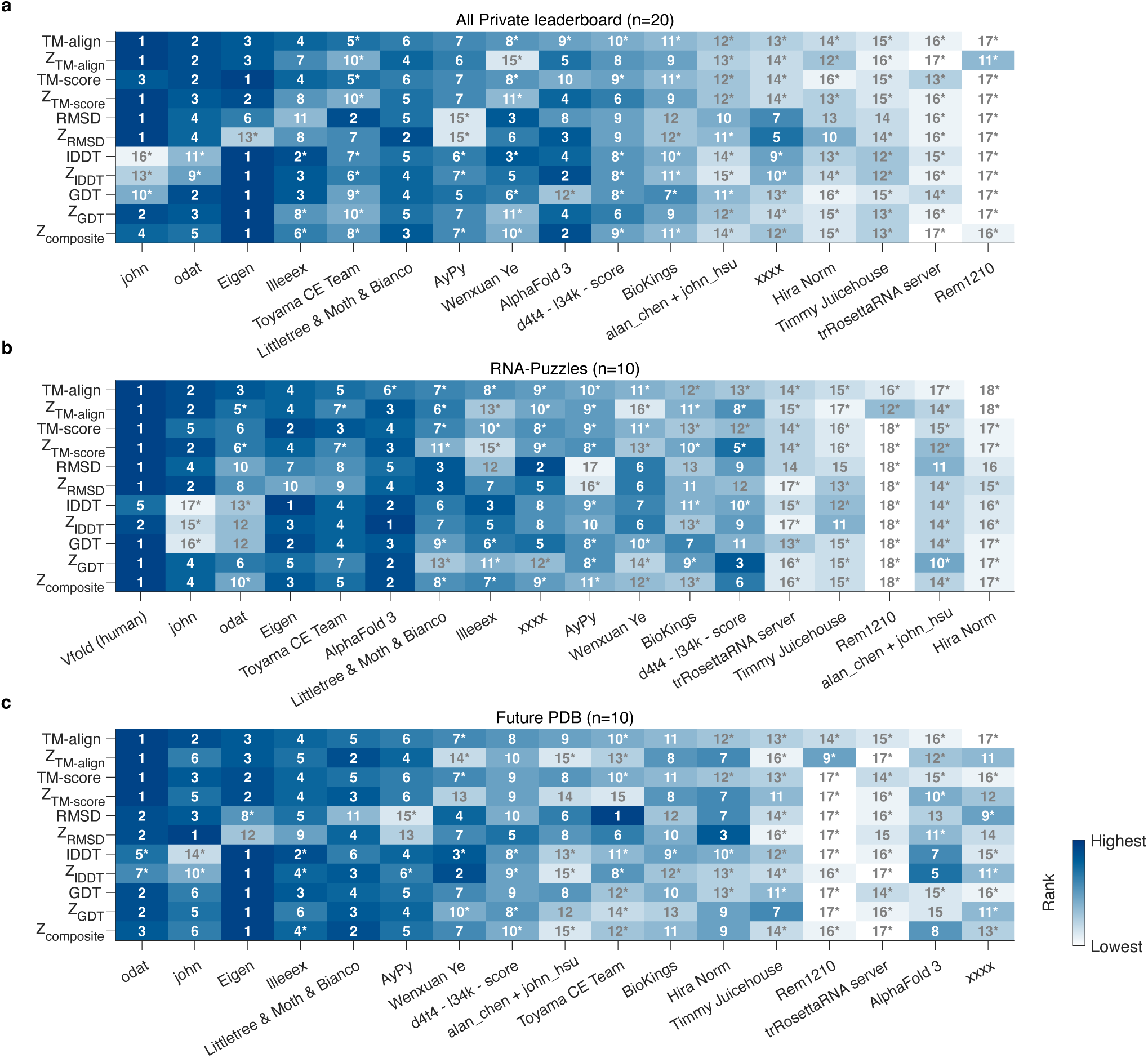
**Group rankings by different metrics and leaderboard subsets**. Rankings of groups based on (a) all 20 Private leaderboard targets, (b) the 10 RNA-Puzzles subset, and (c) the 10 post-competition PDB targets. Vfold ranks appear only in (b), since Vfold did not predict all of (a) or (c). TM-align and TM-score differ only in whether sequence correspondence is forced (see **Methods**); all metrics use C1′ atoms only, for both reference structures and predictions. Z-scores are computed as in **Methods**. Coloring is by rank. For each target set (a-c), john, odat, or Eigen achieved 1st ranking for at least one of the metrics. Asterisks (‘*’) mark groups whose performance was significantly worse (p < 0.05, one-sided paired *t*-test; all *p* values increase with multiple hypothesis testing) than the 1st ranked group in the same row. For TM-align, TM-score, and RMSD, models from odat, john, Eigen and, where available, Vfold are statistically indistinguishable. For GDT, a metric with strong discrimination for small targets, and local accuracy metric lDDT, models from Eigen are the 1st group or are statistically indistinguishable from it.

**Extended Data Fig. 2.**
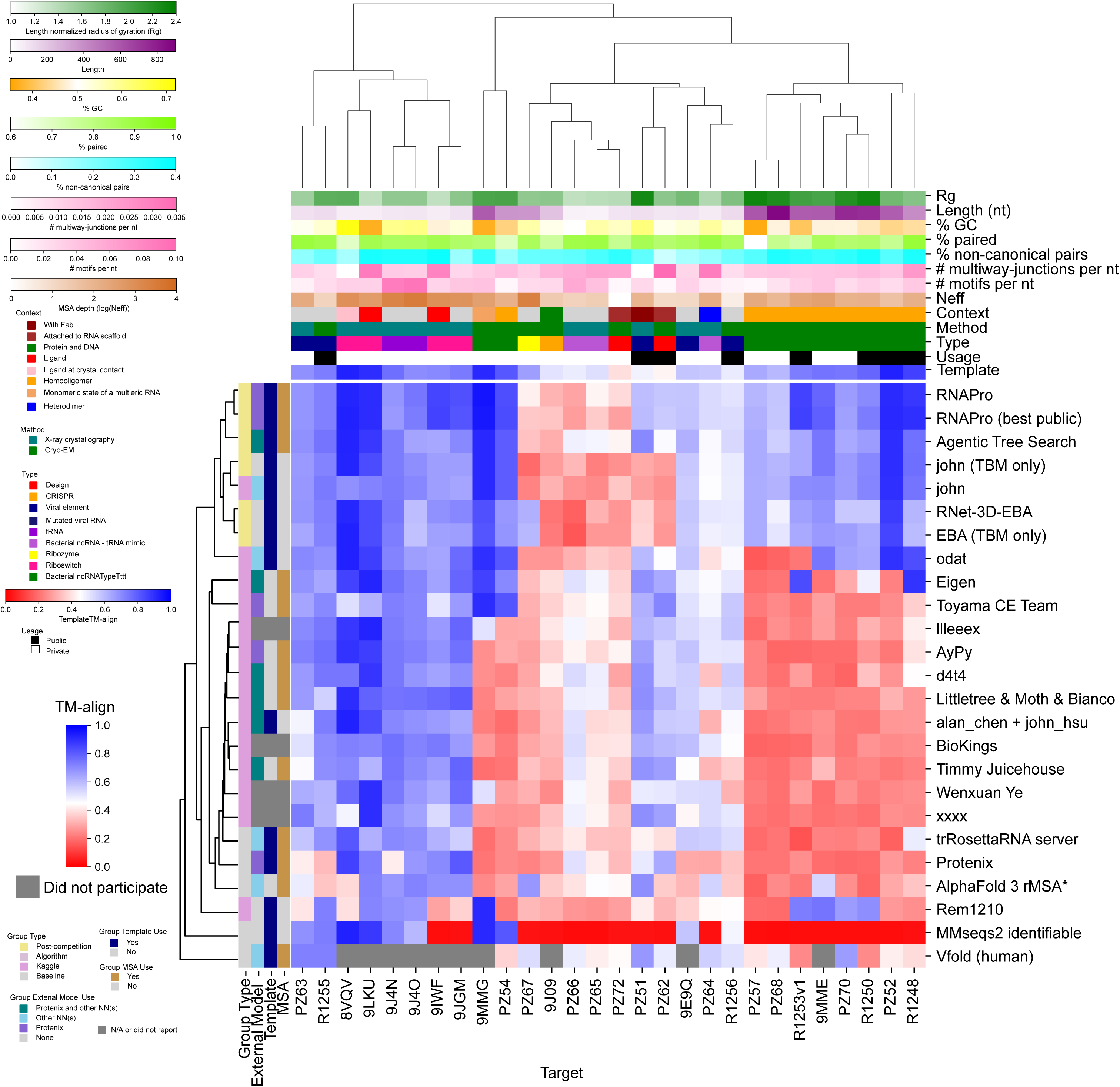
Clustergram of target and group accuracies and features. The best predicted TM-align for groups and targets are plotted. The targets are clustered using Euclidean distance of TM-align accuracy scores. For groups, a tree was created using Euclidean distance as well, but the leaf order was optimized to order the groups’ performance (based on sum of TM-align). The top group was placed at the top, and the tree grew downwards from there. At each node, the branch with the highest average score for its top three teams was added first. If a group did not include models for a target, the box is colored dark gray. The groups and targets are categorized on the left and top of the figure, respectively.

**Extended Data Fig. 3.**
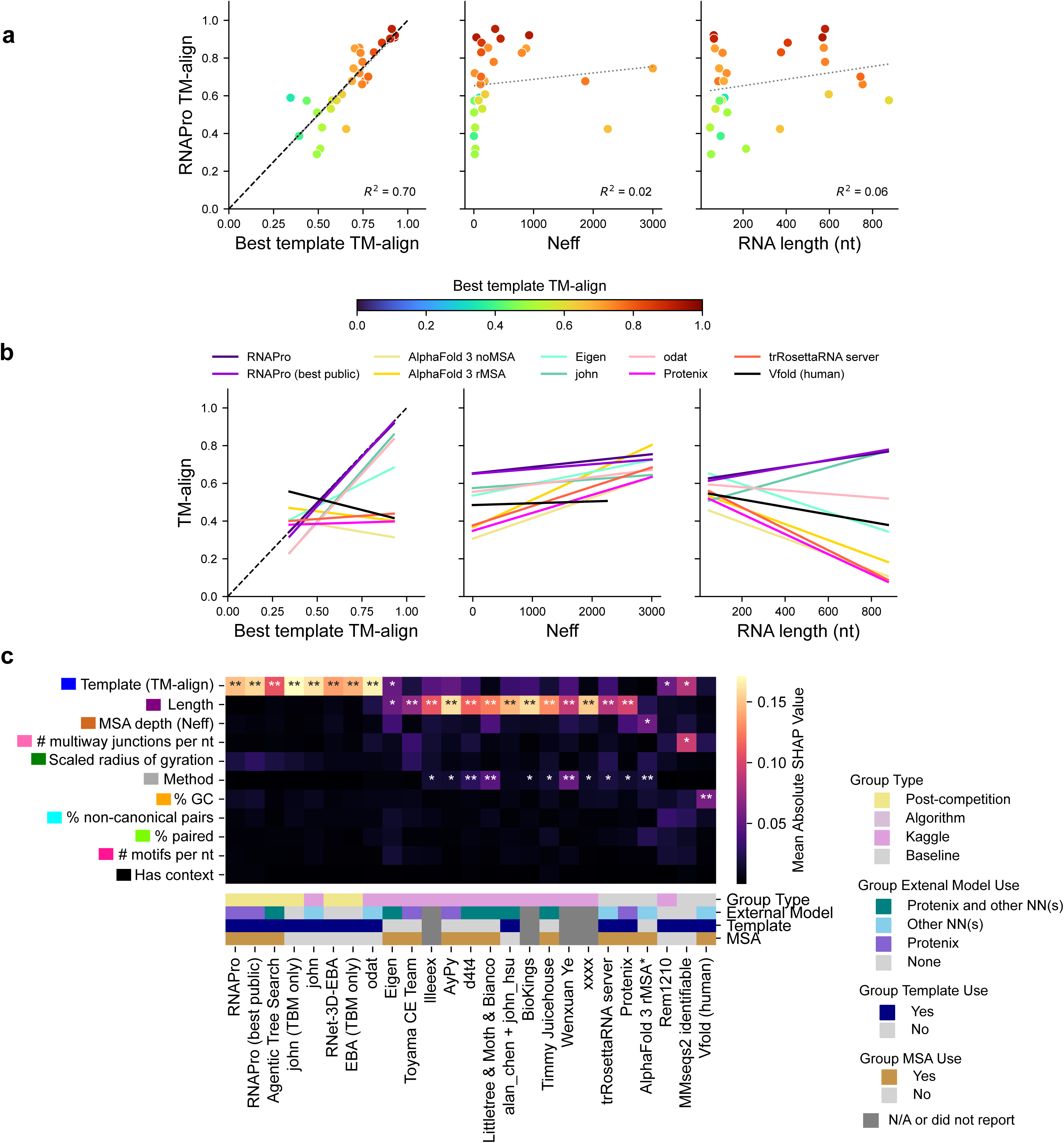
Importance of target properties for each model’s prediction accuracy assessed by SHAP (Shapley Additive exPlanations) analysis. (a) RNAPro best TM-align for each target is plotted against target properties: TM-align of the best template, the normalized number of effective sequences (Neff), and RNA length. The line of best fit is drawn in light grey and the R^2^ is reported at the bottom. The line of best fit for select prediction groups is shown in (b). (c) The mean absolute SHAP value over the 28 targets for which the group participated is plotted from high (yellow) to low (black) for each prediction group and target property. Some prediction groups’ performance is well explained by target template availability, while other groups performance is well explained by target length. Some prediction groups show different trends, e.g., AlphaFold 3’s highest absolute SHAP value is associated with MSA depth. To assess the robustness of feature importance rankings, permutation over target labels (*N*=200) gave a null distribution for SHAP values, and a *p* value was calculated as the number of resulting permutations with mean absolute SHAP values greater than the value calculated from the real data, divided by *N*-1 (*, *p*<0.05; **, *p* <0.01), a one-sided test. All *p* values would increase with corrections for multiple hypothesis testing, not carried out here. SHAP values were calculated with n_estimators=100 (sklearn randomforest), and with the shap Python library.

**Extended Data Fig. 4.**
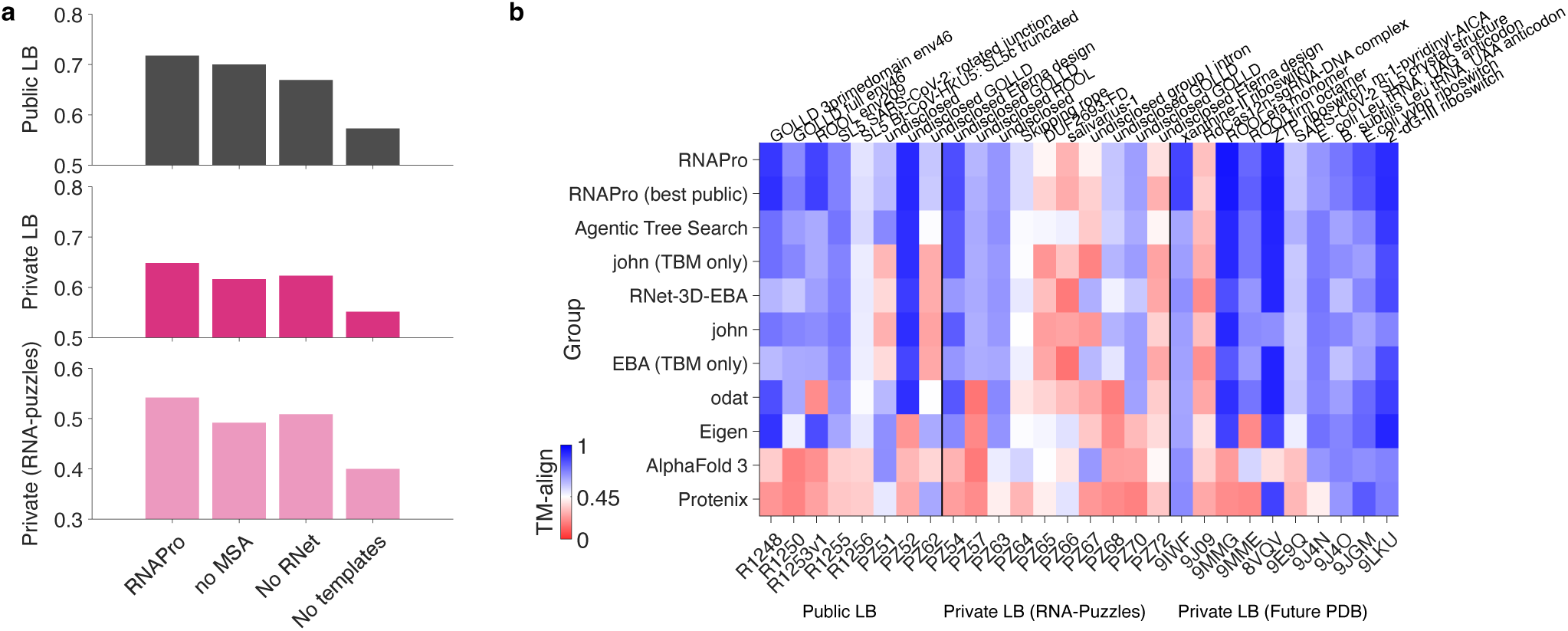
(**a**) Ablation tests confirm the importance of templates relative to other input features for the RNAPro model. (**b**) TM-align scores for individual RNA targets in the final Private leaderboard from post-competition submissions compared to top Kaggle submissions, AlphaFold 3, and Protenix.

**Extended Data Fig. 5.**
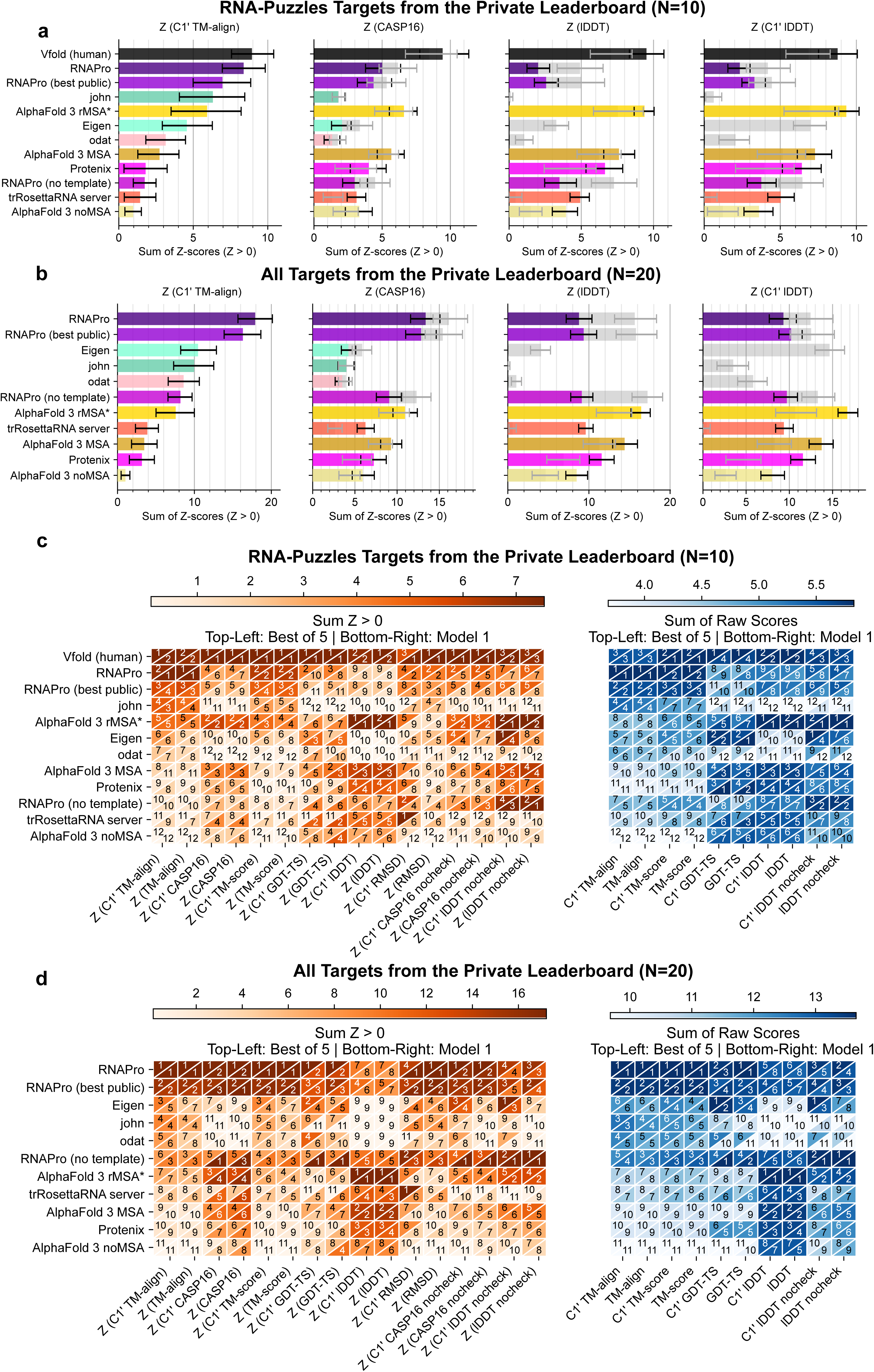
(**a**-**b**) Z-scores for predictors with all-atom models and for C1′-only models from top 3 Kaggle teams with all atoms imputed by Arena for **(a)** the 10 RNA-Puzzle targets from the Private Leaderboard and **(b)** all 20 targets from the Private Leaderboard. Bars show the sum of Z-scores greater than zero across targets for each group and metric; error bars are 68% bootstrap intervals (16th to 84th percentile, ∼1 standard error) from 1,000 bootstrap resamples of the target set. Gray bars are calculations without stereochemical checks for lDDT. (**c-d**) Rankings for **(c)** RNA-Puzzle targets and **(d)** all Private Leaderboard targets with all tested score metrics. Since trRosettaRNA server provided only one model, the top-left corner labels the ranking if only model 1 is used from each group. The bottom-right corner selects the best score amongst the 5 submitted models. Values labeled with “C1′” used C1′ atoms; otherwise, the default atoms for that metric were used (**Methods**). In all panels, the groups are ordered by the sum of Z > 0 for C1′ TM-align. The difference in relative performance of template-using RNAPro and template-free AlphaFold3 between the RNA-Puzzle targets and all Private Leaderboard targets can be explained by the quality of templates available for these targets (mean template TM-align of 0.61 ± 0.04 vs. 0.75 ± 0.05 for RNA-Puzzle vs. other Private Leaderboard targets, respectively).

